# Learning of the same task subserved by substantially different mechanisms between patients with Body Dysmorphic Disorder and healthy individuals

**DOI:** 10.1101/2023.12.19.571882

**Authors:** Zhiyan Wang, Qingleng Tan, Sebastian M. Frank, Yuka Sasaki, David Sheinberg, Katharine A. Phillips, Takeo Watanabe

## Abstract

It is generally believed that learning of a perceptual task involving low-level neuronal mechanisms is similar between individuals. However, it is unclear whether this assumption also applies to individuals with psychiatric disorders that are known to have altered brain activation during visual processing. We investigated this question in patients with body dysmorphic disorder (BDD), a psychiatric disorder characterized by distressing or impairing preoccupation with nonexistent or slight defects in one’s physical appearance, and in healthy controls. Participants completed six training sessions on separate days on a visual detection task for human faces with low spatial frequency (LSF) components. Brain activation during task performance was measured with functional magnetic resonance imaging (fMRI) on separate days prior to and after training. The behavioral results showed that both groups of participants improved on the visual detection task to a similar extent through training. Despite this similarity in behavioral improvement, neuronal changes in the Fusiform Face Area (FFA), a core cortical region involved in face processing, with training were substantially different between groups. First, activation in the right FFA for LSF faces relative to High Spatial Frequency (HSF) faces that were used as an untrained control increased after training in BDD patients but decreased in controls. Second, resting state functional connectivity between left and right FFAs decreased after training in BDD patients but increased in controls. Contrary to the assumption that learning of a perceptual task is subserved by the same neuronal mechanisms across individuals, our results indicate that the neuronal mechanisms involved in learning of a face detection task differ fundamentally between patients with BDD and healthy individuals. The involvement of different neuronal mechanisms for learning of even simple perceptual tasks in patients with BDD might reflect the brain’s adaptations to altered functions imposed by the psychiatric disorder.

## Introduction

Most empirical and computational studies assume that mechanisms of learning of the same perceptual task are similar across individuals (Plomin, 2001; Yang et al., 2020). This leads to the prediction that individuals with altered brain function due to psychiatric disorder should learn by employing, perhaps to an altered extent, similar neural mechanisms as healthy individuals. To test these assumptions, we compared visual perceptual learning (VPL) of faces and neuronal mechanisms of learning involved between patients with body dysmorphic disorder (BDD) and healthy individuals. VPL refers to a performance improvement in a visual task as a result of visual experience or training and is regarded as a manifestation of visual plasticity (Watanabe et al., 2001; Watanabe et al., 2002; Seitz and Watanabe, 2003; Watanabe, Nanez and Sasaki, 2010; Shibata et al., 2011; Watanabe and Sasaki, 2015; Shibata et al., 2017; Bang et al, 2018; Tamaki et al., 2020; Frank et al., 2021; Lu and Dosher, 2022; Zhang, et al., 2023; Klorfeld-Auslender et al., 2022; Seitz, 2021; Hung and Carrasco, 2023; Harris et al, 2015). BDD is a relatively common psychiatric disorder characterized by distressing or impairing preoccupation with nonexistent or slight defects in one’s appearance, most often the face (Simmons and Phillips, 2017). BDD is associated with marked impairment in functioning, poor quality of life, and high rates of suicidality (Phillips et al., 2005; Snorrason et al., 2019). Previous results showed that BDD patients exhibited perceptual and neuronal differences in face processing from healthy controls. First, there are perceptual differences for holistic processing of faces corresponding to low-frequency (global) components between BDD patients and healthy controls (Feusner et al., 2010; Jefferies et al., 2012; Mundy et al., 2014). For example, BDD patients showed less of an inversion effect compared to healthy controls in the recognition of an inverted face (Feusner et al., 2010). It is believed that such faces are recognized more slowly and less accurately by healthy controls as a result of disrupted holistic processing in an upside-down face (Yin, 1969; Farah, Tanaka & Drain, 1995; Leder & Bruce, 2000; Valentine, 1988). Second, compared with healthy controls, BDD patients showed weaker functional magnetic resonance imaging (fMRI) activation in the left occipital and fusiform cortices including the Fusiform Face Area (FFA) during processing of low frequency faces (Feusner, Moody et al., 2010; Li et al., 2015). The FFA is a core brain region for face processing (Kanwisher, McDermott et al., 1997; Rossion, Gauthier et al., 2000; Grill-Spector, Knouf et al., 2004; Kanwisher and Yovel, 2006; Rotshtein, Vuilleumier et al., 2007; Goffaux, Peters et al., 2011; Haxby and Gobbini, 2011; Richler and Gauthier, 2014; Duchaine and Yovel, 2015; Yovel, 2016). Third, compared with healthy controls, BDD patients exhibited altered connectivity and information transfer between face processing regions in different hemispheres (Moody, Sasaki et al., 2015). Together, these findings suggest that there are substantial differences in the perception of faces and in the neuronal circuits in BDD.

Although these studies suggest that BDD patients process low frequency faces differently than healthy participants (Feusner, Townsend et al., 2007; Feusner, Moody et al., 2010), processes involved in the development of learning in BDD patients are unclear. To address this, we trained BDD patients and healthy controls on an identical visual detection task, using faces consisting solely of low spatial frequency (LSF) components. Before the first and after the final training sessions, participants’ neural responses during a face detection task were assessed using fMRI. Blood oxygen level dependent (BOLD) responses for FFA were calculated and compared between BDD patients and healthy controls before and after training. Furthermore, we also measured resting state functional connectivity between left and right FFA.

The results of the present study showed that training on LSF faces significantly improved only the detectability of LSF faces to a similar extent in each participant group. Despite this similarity in learning between the groups, changes of BOLD responses (i.e., activity increase or decrease with learning) and functional connectivity between the right FFA and the left and right FFA (i.e., connectivity increase or decrease with learning) occurred in opposite directions (activity or connectivity increase vs. decrease) between BDD patients and healthy controls.

These results challenge the assumption that learning of a perceptual task and the resulting pattern of performance improvement are driven by the same underlying mechanisms across individuals, irrespective of the presence or absence of a psychiatric disorder. Instead, the findings indicate that individuals with a psychiatric disorder can achieve comparable or even greater performance improvements in a perceptual learning task with training as healthy participants but employ different neuronal mechanisms for this learning. However, they appear to rely on different strategies and neuronal mechanisms compared to those with normal brain function when adapting to a new environment, making the best use of their present abnormal brain mechanisms.

## Methods

### Participants

Study participants were 9 individuals with BDD (22 - 52 years old, mean age = 34.9 +/- SD 11.34 years, 8 female and 1 male) recruited from the Rhode Island Hospital and 10 healthy control participants of a similar age range (20 – 60 years old, mean age = 27.8 +/- SD 11.81 years, 5 female and 5 male) recruited from the Brown University community. Study inclusion criteria for the BDD group were having BDD as the primary diagnosis for >6 months and a score >24 on the Yale-Brown Obsessive-Compulsive Scale Modified for BDD, indicating BDD of at least moderate severity (Phillips et al, 1997). BDD was diagnosed by a psychiatrist with expertise in BDD (K.A.P.) using the Structured Clinical Interview for DSM-IV-Patient Version (SCID-I/P) (First et al., 1995). Exclusion criteria for both groups included 1) Current manic episode, history of a psychotic disorder, or a substance use disorder in the past 3 months (determined with the SCID-I/P); 2) a positive urine drug screen for a drug that was not prescribed; 3) Receipt or initiation of cognitive-behavioral therapy for BDD during the study; 4) Initiation of psychotropic medication or a dose change during the study; medication could be taken if the dose had been stable for at least 2 months before study participation. Healthy controls could not have a psychiatric disorder (determined by the SCID-I/P) or be taking psychoactive medication. All participants had normal or corrected-to-normal vision. They gave written informed consent prior to participation. The study was approved by the institutional review boards of Brown University and the Rhode Island Hospital.

### Experiment Design

The study consisted of 4 stages as shown in Fig. 1A: the sensitivity measurement stage (1 session), pretest stage (1 session), training stage (6 sessions) and posttest stage (1 session). Each session was scheduled at least one day apart. The sensitivity measurement and training stages were conducted in a dimly lighted psychophysical testing room outside the MRI scanner. The pretest and posttest stages were conducted inside the MRI scanner (see below).

**Figure 1.**
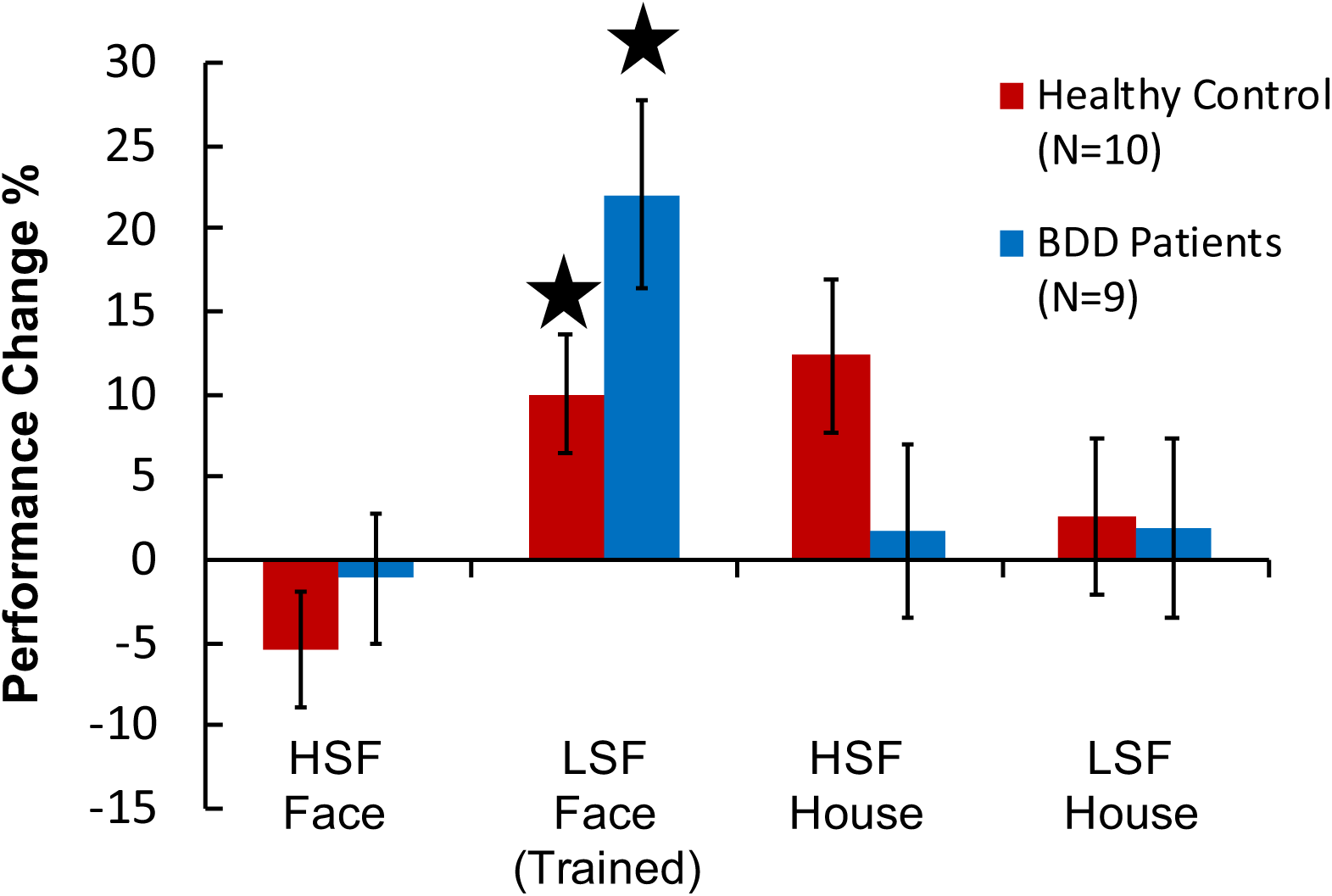
Behavioral results. Mean ± standard-error-of-the-mean (SEM) performance change across patients with body dysmorphic disorder (BDD) and healthy controls. Zero on the y-axis corresponds to performance before training. Values above zero on the y-axis indicate that performance improved after training. The results showed that both groups improved on the detection of LSF faces through training. The improvement did not transfer to other untrained conditions. * *p* < 0.05.

### Stimuli

For the sensitivity measurement, pretest and posttest stages, a set of face images (50% females and 50% males) with neutral facial expressions (Psychological Image Collection at Sterling, https://pics.stir.ac.uk) were used. In addition, house images (Filliter et al., 2016) were used as a control for identifying changes that are specifically related to LSF faces (Fig. 1B). There were 4 different stimulus categories of images. The LSF faces and LSF houses consisted of the frequency components of images that were less than 10 cycle/degree. The HSF faces and HSF houses consisted of the frequency components of images that were larger than 15 cycle/degree. Four different face/house images were used for each category. During the training stage, only LSF faces were used. Twelve different face images were used during the training stage. For training, a different set of face images from those for the sensitivity measurement and testing stages were used to ensure that training results did not originate from learning of specific face images but learning of LSF components of faces in general. The images were in a gray scale and all images had the same mean contrast. Each face and house image was tilted 45^0^ away from the upright position to make the task challenging for participants. Visual noise was embedded in an image by replacing a subset of pixels of the image with values selected from a Gaussian distribution. At the center of the display, a white bull’s eye on a gray disk with a radius of 0.75 deg was presented. The faces and houses were presented in a square of 3 x 3 arc deg on a homogeneous gray square background of 4 x 4 arc deg.

In addition, gray-scale images of faces and places were used to define regions-of-interest (ROIs) in each participant (see *ROI Definitions* below) following the approach of previous studies (e.g., Polk et al., 2007). These images consisted of upright faces, phase scrambled faces (spatial frequency was protected), upright houses and phase scrambled houses. The phase spectra of the face and house images were obtained and randomized using fast-Fourier transformation. Phase-scrambled face and house images were then created with inverse Fourier transformation. The size of these images was the same as of the stimuli used for training.

### Sensitivity Measurement

Each participant’s thresholds for detecting face and house stimuli with different spatial frequencies were measured using a two-interval-forced-choice (2IFC) task. Each face and house image with each spatial frequency was tested in a different block. Each trial contained two 200 ms stimulus intervals with a 600 ms blank interval in between. The trial ended after a 1000 ms interval during which the participant was asked to make a response. During one of the two stimulus intervals, a face or house image embedded in Gaussian noise was presented. In the other stimulus interval, an image consisting only of noise was presented, as shown in Figure 1C. Participants were asked to indicate whether the first or second stimulus interval contained a face or house by pressing one of two keys on the computer keyboard. The signal-to-noise (S/N) level in a stimulus was controlled using a 3-down-1-up staircase procedure, approximating a response accuracy of 84% correct. The initial S/N ratio was 25%. The step size of the staircase was 0.05 log units. After 10 staircase reversals, the testing block terminated. The S/N ratio threshold was calculated as the geometric mean of the last six reversals.

### Pretest and Posttest

The pretest and posttest stages consisted of several substages, which were all conducted inside the MRI scanner: structural MRI acquisition, functional MRI acquisition during task performance, localization of ROIs and functional connectivity measurement. First, during the structural MRI acquisition, a high-resolution anatomical scan of each participant’s brain was collected using a T1-weighted MPRAGE sequence [256 sagittal slices, voxel size = 1 x 1 x 1 mm, time-to-repeat (TR) = 1980 ms, time-to-echo (TE) = 3 ms, flip angle = 9°]. Second, during functional MRI acquisition, participants performed the 2IFC task on face and house stimuli with LSF and HSF components while BOLD responses were measured using a T2*-weighted echoplanar imaging (EPI) sequence (33 transverse slices, voxel size = 3 x 3 x 3 mm, TR = 2000 ms, TE = 25 ms, flip angle = 90°). 6 such fMRI runs were conducted using a block design in which blocks of stimulation were followed by baseline blocks without stimulation. Throughout each fMRI run, participants were instructed to maintain fixation at a white bull’s eye at the center of the display. Each fMRI run lasted 150 s and consisted of twelve 12 s-long blocks. Each fMRI run commenced with a 4 s-long fixation period and terminated with a 2 s-long fixation period. Each block consisted of 6 trials with images from only one category. Each trial lasted 2 s and consisted of two 200 ms stimulus intervals with a 600 ms interval in between, followed by a 1000 ms response interval after the second stimulus interval. The S/N ratio level for each stimulus category was set to each participant’s threshold measured during the sensitivity measurement stage. Third, the localization of FFA was conducted with 2 fMRI runs. During these fMRI runs images of faces and places were presented in blocks of 12 s each. Each block consisted of one type of the four image categories (i.e., upright faces, upright houses, phase scrambled faces and phase scrambled houses). Participants were instructed to maintain fixation at a white bull’s eye at the center of the display and perform a 1-back task. In this task they were asked to press a button when the currently presented face or house image matched the image presented immediately before the current image, which occurred occasionally and in an unpredictable fashion (Berman et al., 2010). Each fMRI localizer run consisted of 16 blocks and lasted 200 s. Each run started and terminated with a 4 s-long fixation period. The same EPI sequence parameters as in the functional task runs were used. Fourth, resting state functional connectivity was measured using the same EPI sequence parameters as in the functional task runs. We investigated functional connectivity based on the resting state time-course of the BOLD signal (Davies-Thompson & Andrews, 2012; Norman-Haignere et al., 2012) between left and right FFA before and after training. The resting state BOLD signal was measured in a 410 s-long fMRI run during which participants were instructed to maintain fixation at a fixation cross at the center of the display with no other face or house stimuli presented and no other task. The resting state runs were conducted at the beginning of the pretest and posttest stage to avoid artefacts from tiredness and influence from other fMRI runs.

### Training

The training stage was conducted in a dimly lighted psychophysical room over the course of 6 sessions on separate days. Participants performed the same 2IFC task as during the sensitivity measurement stage but using only face images with LSF components. Each session consisted of 10 blocks. Each block was conducted with a 3-down-1-up staircase procedure. The initial S/N ratio was set to 25%. The step size of the staircase was 0.05 log units. After 10 staircase reversals, the training block terminated. The S/N ratio threshold of each block was defined as the geometric mean of the last six reversals. Participants performed approximately 500 trials for each session, resulting in a session duration of 30 min.

### Apparatus

For training visual stimuli were presented on an LCD display (1920 X 1200 pixel resolution, 60 Hz refresh rate, viewing distance 50 cm). Participants were seated in front of a computer screen and used a chinrest. They were provided a keyboard for response. Visual stimuli were presented in the scanner via a projector and participants viewed the stimuli using a headcoil mounted mirror. Visual stimulation in training and scanning was controlled via Psychtoolbox (Brainard, 1997; Pelli, 1997) running in MATLAB (The Mathworks; Version R2015b). Scanning was performed using a 3T Siemens Prisma MRI scanner and a 64-channel head/neck coil. Participants were given an MRI-safe button box for response.

### fMRI Data Analysis

#### Preprocessing

Structural and functional MRI scans were preprocessed and analyzed using the FreeSurfer software package (Version 4.5) and the FSFast toolbox (Dale et al., 1999, Fischl et al., 1999). Each participant’s high-resolution anatomical scan was reconstructed and inflated. fMRI images from each run were motion-corrected, realigned to the first volume from the first run of each session as a template, coregistered to the high-resolution reconstructed scan of the brain (collected in the pretest stage), spatially smoothed using a three-dimensional Gaussian kernel (FWHM = 5mm), intensity normalized and slice time corrected.

#### Definition of FFA

To obtain BOLD activation for the localization of FFA, a general linear model (GLM) assuming a gamma shaped hemodynamic response function (HRF) with a delay of 2.25 s and a dispersion of 1.25 s was fitted to the BOLD signal measured in the fMRI localizer runs. The model contained four regressors-of-interest for four stimulus conditions: upright faces, scrambled faces, upright houses, scrambled houses. The model was also fitted with nuisance regressors including motion correction parameters, polynomial drift regressors and temporal whitening. We localized FFA in the fusiform gyrus by using the contrast of significantly stronger BOLD activation for faces than scrambled faces in the functional localizer scan (*p* < 0.001, uncorrected). The center of gravity and the cluster size of FFA were similar between BDD patients and healthy controls (see Table 1).

**Table 1.** Mean MNI coordinates (mm) with standard deviation (mm) of the center of gravity of FFA across participants. Cluster size corresponds to number of voxels in functional space.

|  |  | **X** | **Y** | **Z** | **cluster size**<br>**(voxels)** |
| --- | --- | --- | --- | --- | --- |
| **BDD** | Left FFA | - 40 ± 1.4 | - 50 ± 8.2 | -16 ± 2.0 | 272 ± 202 |
|  | Right FFA | 42 ± 2.0 | - 55 ± 6.5 | -14 ± 2.1 | 231 ± 193 |
| **Controls** | Left FFA | - 40 ± 1.1 | - 52 ± 7.2 | -16 ± 1.3 | 211 ± 177 |
|  | Right FFA | 40 ± 1.7 | - 51 ± 4.7 | -15 ± 1.6 | 300 ± 118 |

#### fMRI Activation During 2IFC Task Performance

Another GLM using the same gamma shaped HRF function and nuisance regressors as in the GLM for ROI localization was carried out to calculate BOLD activation while participants performed the 2IFC task on face and house stimuli with LSF and HSF components. The GLM included four regressors-of-interest for four stimulus categories: LSF faces, HSF faces, LSF houses and HSF houses. Beta coefficients, reflecting the BOLD response, for each condition were extracted for FFA.

#### Functional Connectivity Between Left and Right FFA

Functional connectivity using resting state BOLD measurements was calculated following the same approach as in a previous study (Shmuel et al., 2021). The time-course signals of cerebrospinal fluid and white matter were extracted from the preprocessed functional resting state scans using principal component analysis. These time-courses together with motion correction parameters and linear scanner drift were used as nuisance regressors for a GLM of the resting state BOLD measurements to account for and remove variance-of-no-interest. The residual BOLD signals of the fitted GLM corresponding to the resting state BOLD time-course corrected for variance-of-no-interest were extracted from voxels in left and right FFA and averaged across voxels within the ROIs. Functional connectivity was calculated between resting state BOLD signals in the left and right FFA as Fisher-transformed Pearson correlation coefficient.

#### Statistics

Normality of the data was tested with the Shapiro-Wilk test. The results showed no violations of the assumption of normality. ANOVAs were used for between and within group comparisons. *T*-tests were conducted for post-hoc analyses. The statistical analyses were carried out using MATLAB (Version R2019b) and SPSS (IBM; Version 24).

## Results

### Behavioral Results

Performance changes for HSF faces, LSF faces, HSF houses and LSF houses in each participant group were obtained by calculating [(posttest accuracy – pretest accuracy) / pretest accuracy × 100%]. We performed a mixed-design ANOVA with ‘Group’ [BDD vs. control] as the between-subject factor and ‘Condition’ [HSF face, LSF face, HSF house vs. LSF house] as the within-subject factor on performance change. The ANOVA yielded a significant main effect of ‘Condition’ [*F*(3, 51) = 4.547, *p* = 0.007, partial *11^2^* = 0.211], indicating that participants showed greater performance improvements for LSF faces than for other conditions after training (Figure 2). We further conducted post-hoc *t*-tests for each condition separately for BDD patients and healthy controls. There were significant performance improvements for LSF faces after training in the BDD group [one-sample *t*-test of performance change in posttest against zero corresponding to pretest performance; *t*(8) = 3.892, *p* = 0.005, Cohen’s *d* = 1.29] and the control group [*t*(9) = 2.775, *p* = 0.02, Cohen’s *d* = 0.91]. The magnitude of performance improvement for LSF faces did not differ significantly between groups [two-sample *t*-test; *t*(17) = 1.829, *p* = 0.085]. No other significant performance changes in any condition in any participant group were found. Together, these results show that training to detect LSF faces yielded performance improvements indicative of VPL only for the trained stimuli in both BDD patients and healthy controls. The magnitude of performance improvement was similar between patients and controls.

### fMRI Results

#### Activation Changes in FFA Associated with VPL

A 2 x 2 x 2 mixed-design ANOVA with group (BDD vs. controls) as a between-subject factor and hemisphere (left vs. right) and test (pretest vs. posttest) as within-subject factors was applied to beta difference scores between LFS and HFS faces in FFA. There was a significant three-way interaction [*F*(1, 17) = 15.936, *p* = 0.001, partial *11^2^* = 0.484], indicating that activation changes from pretest to posttest differed between left and right FFA and between participant groups (Figure 3).

**Figure 3.**
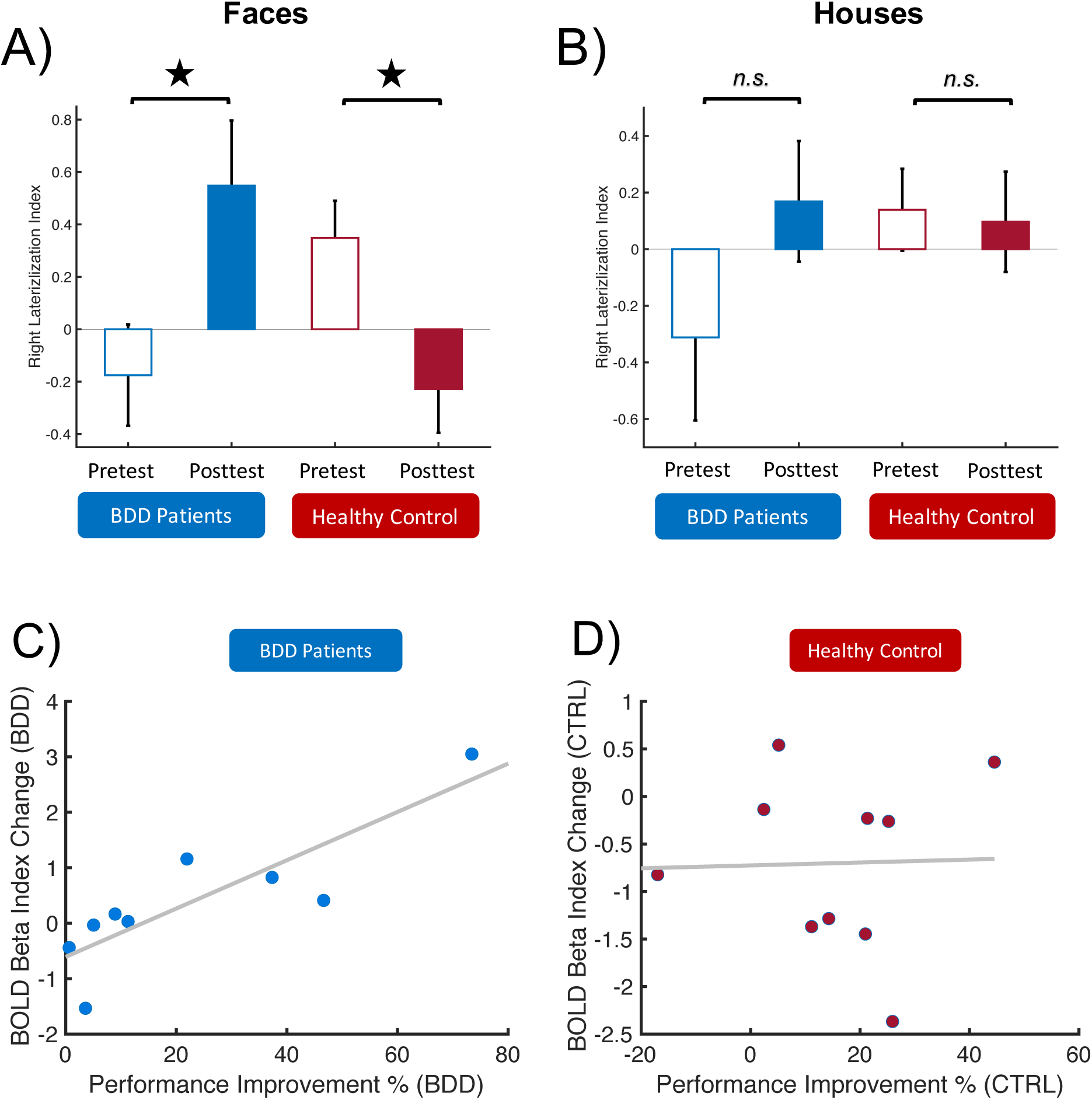
fMRI results. A) Mean ± SEM BOLD beta index of FFA right lateralization for comparison between LSF and HSF faces across BDD patients and healthy controls. Greater values on the y-axis correspond to greater FFA right lateralization. B) Same as A) but for the comparison between LSF and HSF houses. C) Correlation between performance improvement and BOLD beta index of FFA right lateralization across BDD patients. Each dot shows the result from a different participant. D) Same as C) but across healthy controls. * *p* < 0.05.

To measure the three-way interaction between the factors hemisphere, test, and group, we calculated a right lateralization index in FFA based on the BOLD response for trained face and untrained house, respectively. The right lateralization index for face stimuli was defined as the subtraction of the trained face response in the left FFA [beta(LSF)-beta(HSF)] from the trained face response in the right FFA [beta(LSF)-beta(HSF)]. The right lateralization index for house stimuli was defined as the subtraction of the house response in the left FFA [beta(LSF)-beta(HSF)] from the house response in the right FFA [beta(LSF)-beta(HSF)]. The right lateralization indices for each participant group and test stage are shown in Figure 3A and B for the face and house stimuli, respectively. We applied a 2 x 2 mixed design ANOVA with group (BDD vs. controls) as the between-subject factor and test (pretest vs. posttest) as the within-subject factor to right lateralization indices separately for faces and houses (Figure 3A and B). For face stimuli (Figure 3A), there was a significant interaction effect of group and test [*F*(1, 17) = 15.936, *p* = 0.001, partial *11^2^* = 0.484]. Post-hoc *t*-tests showed that the right lateralization index in the BDD group significantly increased from pretest to posttest [*t*(8) = 3.009, *p* = 0.017, Cohen’s *d* = 1.003], whereas it significantly decreased in the control group from pretest to posttest [*t*(9) = **-**2.6137, *p* = 0.028, Cohen’s *d* = 0.827]. For the house stimuli (Figure 3B), there was no significant interaction between group and test [*F*(1,17) = 2.042, *p* = 0.17], no main effect of group [*F*(1,17) = 0.550, *p* = 0.468] and no main effect of test [*F*(1,17) = 1.434, *p* = 0.248].

To test whether performance improvements were associated with changes in the right lateralization index from pretest to posttest, we calculated a Pearson correlation coefficient. We found a significant correlation in BDD patients [*r* = 0.780, *p* = 0.013] such that patients with greater behavioral improvements tended to show more pronounced increases of right lateralization. No such significant correlation was found in controls [*r* = 0.121, *p* = 0.740] (see Figure 3C and D). Note that there were no outliers in the change of lateralization index or performance improvement in either group (Grubb’s test, *p* = 0.382 for BDD patients and *p* = 0.706 for healthy controls).

#### Functional Connectivity Between Left and Right FFA

A 2 x 2 mixed design ANOVA with group (BDD vs. controls) as a between-subject factor and test (pretest vs. posttest) as a within-subject factor on functional connectivity between left and right FFA showed a significant interaction between group and test [*F*(1, 17) = 18.75, *p* = 0.0005, partial *11^2^* = 0.524], indicating that connectivity changed differently between BDD patients and controls from pretest to posttest (Figure 4). Post-hoc *t*-tests showed a significant decrease in functional connectivity between left and right FFA in BDD patients from pretest to posttest [paired-sample *t*-test; *t*(8) = −4.221, *p* = 0.003, Cohen’s *d* = 1.407] and a significant increase in controls [*t*(9) = 2.698, *p* = 0.025, Cohen’s *d* = 0.853].

**Figure 4.**
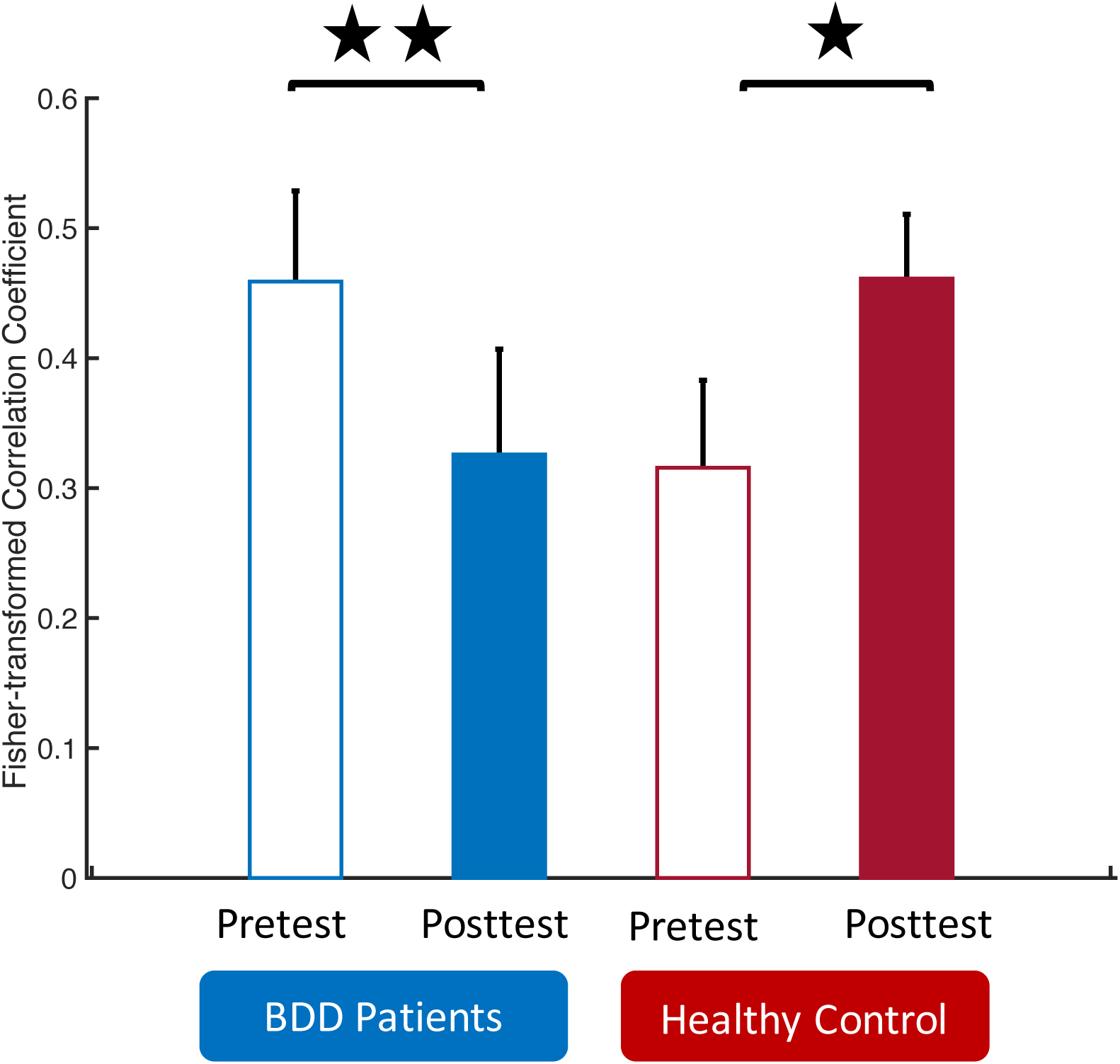
Mean ± SEM change in Fisher-transformed correlation coefficient between left and right FFA across BDD patients and healthy controls. Greater values on the y-axis correspond to greater connectivity. ** *p* < 0.01. * *p* < 0.05.

## Discussion

In this study, we examined whether VPL of faces occurs in a similar fashion between BDD patients and healthy control participants. To achieve this, we conducted visual detection training for LSF faces in BDD patients and controls. To our knowledge, this is the first study to examine neuronal changes in BDD patients in association with perceptual training. Both groups exhibited similar improvements in performance through training. However, the neural changes underlying this learning process, reflecting neuronal mechanisms involved, showed striking differences between the groups. In the right FFA, a core cortical region involved in face processing, activation for LSF faces relative to HSF faces increased after training in BDD patients, while it decreased in controls. Moreover, there were notable differences in resting state functional connectivity between left and right FFA, such that connectivity decreased in BDD patients but increased in controls through training.

These findings suggest that comparable improvements in performance can be achieved despite significant differences in the patterns of neuronal changes in FFA in BDD patients and controls. Consequently, the results suggest that learning observed in BDD patients does not necessarily involve the same underlying neuronal mechanisms as in healthy controls. This may reflect that BDD patients may adopt distinct strategies to optimize neuronal resources for adapting to new environments, thereby leveraging their current brain mechanisms in unique ways.

Our findings have potential clinical implications. Although serotonin-reuptake inhibitor medications and cognitive behavioral therapy are often effective treatments for BDD (Philips 2017; Wilhelm et al., 2019), not all patients improve, and additional BDD treatments are needed. There are no established perceptual training paradigms for BDD based on neuronal mechanisms, despite increasing evidence of perceptual abnormalities in BDD. One recent study suggests that enhancing functional connectivity in visual areas responsible for holistic processing may increase the affective evaluation of patients’ body evaluation (Wong et al, 2021). Our study similarly suggests that our training paradigm, which increased sensitivity for low spatial frequencies and thus has the potential to enhance holistic visual processing, has promise as a potential treatment for BDD. Moreover, previous studies have suggested that BDD patients have a dominance of brain volumes and activities in the left hemisphere (Feusner et al., 2007; Feusner et al., 2023). In our study such left hemisphere dominance when viewing LSF faces in BDD patients decreased after training and their brain activities became similar to that of the controls before training. However, due to the relatively small sample size and a dominance of female BDD patients, further study is needed to clarify the clinical effects.

Although not significant, there was a slight behavioral improvement for the HSF house condition in the control group. It is possible that the neuronal changes associated with training on LSF faces could also have an effect on the detection of HSF houses. However, further study is necessary to explore if there is an interaction between VPL of LSF face and HSF house.

To conclude, these results shed light on the complexity of brain function and the diversity of adaptations that can occur in response to learning tasks. They also have potential clinical implications. Further research is required to delve deeper into the specific mechanisms responsible for these differences, providing a comprehensive understanding of the neural basis of learning and its implications for BDD and perhaps other psychiatric disorders involving deficits in holistic visual processing such as anorexia nervosa (Li et al., 2015).

## Acknowledgment

This study was funded by a Dean’s Emerging Areas of New Science (DEANS) Award from the Brown University Division of Biology and Medicine to KP, DS, TW, the National Institutes of Health (NIH) R01EY031705 and R01EY019466 to TW and R01EY027841 to YS.

